# Obscuration to Clarity: Bone Suppression for Enhanced Localization in Pneumothorax Segmentation of Chest Radiographs

**DOI:** 10.64898/2026.02.17.706467

**Authors:** Ananya Shukla, Amog Rao, S. Siddharth, Rina Bao

**Affiliations:** Plaksha University, India; Boston Children’s Hospital & Harvard Medical School, USA

**Keywords:** Bone Suppression, Pneumothorax, Boundary Aware Segmentation, Computer-Aided Diagnosis

## Abstract

Chest radiography (CXR) is a primary modality for assessing cardiopulmonary conditions, but its effectiveness is limited by anatomical obstructions (e.g., ribs, clavicles) that hinder accurate pneumothorax segmentation, boundary delineation, and severity estimation. While deep learning–based bone suppression improves soft-tissue visibility, its utility for precise pixel-wise localization remains underexplored. This study investigates the downstream application of bone suppression for pneumothorax segmentation, integrating it as a preprocessing step to mitigate bony obscuration. We evaluate its impact across CNN and Vision Transformer models on two public datasets, where models trained on bone-suppressed CXRs significantly outperform (*p <* 0.05) non-suppressed counterparts, achieving up to 17% improvement in Mean Average Surface Distance (MASD), 4.9% in Dice Similarity Coefficient (DSC), and 5.9% in Normalized Surface Dice (NSD), alongside a 9.5% gain in Matthew’s Correlation Coefficient (MCC). These results demonstrate bone suppression as an architecture-independent enhancement for pneumothorax localization, improving the reliability of automated CXR interpretation.

## 1. INTRODUCTION

Chest X-Rays (CXRs) are the primary modality for detecting cardiopulmonary abnormalities due to low radiation exposure but their 2D projection causes rib and clavicle overlap, obscuring subtle lung abnormalities and leading to missed diagnoses [1]. Bone suppression [2] computationally removes bony structures to enhance soft-tissue visibility, and recent deep learning–based methods have improved detection of tuberculosis and lung nodules without additional radiation exposure.

Pulmonary conditions such as pneumothorax, caused by air accumulation in the pleural cavity can lead to lung collapse, ranging in severity from asymptomatic to hemodynamic instability or death [3]. Its assessment is particularly challenging, as collapsed lung regions are often obscured by the overlying ribs and clavicles. Given the need for precise localization, our objective is to assess whether segmentation of bone-suppressed CXRs could enable improved severity estimation, establishing their clinical utility in early diagnosis and triaging of critical cases. To achieve this goal we propose

- A novel application of bone suppression for pixel-wise pneumothorax segmentation. Our approach integrates a ResNet-based bone-suppression model as a pre-processing step, reducing anatomical obstructions in CXRs.
- Upon extensive evaluation on two datasets, we demonstrate that bone suppression enhances segmentation performance across architectures, benchmarking it across Convolution Neural Networks (CNNs) and Vision Transformers (ViTs), indicating architecture-independent performance gains from bone-suppressed inputs.
- A shift in focus from classification-centric bone suppression to segmentation tasks, evaluated using boundary-aware metrics such as Mean Average Surface Distance (MASD) for boundary deviation and Normalized Surface Distance (NSD) for boundary accuracy. Crucially, these metrics along with GradCAM [4] activation maps demonstrate the impact of bone suppression on pneumothorax localization and boundary delineation in segmentation.

## 2. RELATED WORK

A widely used approach for reducing anatomical obstructions in CXRs is dual-energy subtraction (DES), which relies on specialized acquisition protocols and lacks portability [10]. To address this limitation, as shown in Table 1, Rajaraman et al. [5] proposed a bone suppression framework to enhance tuberculosis classification using VGG-16, trained on the JSRT dataset. The models significantly improved AUC on the Shenzhen (0.899 to 0.954) and Montgomery (0.857 to 0.964) datasets, highlighting enhanced model sensitivity. Rajaraman [6] later introduced DeBoNet, an ensemble of U-Net and FPN. The model is further adapted for COVID-19 classification, outperforming non-suppressed CXRs with an AUROC of 0.99. Recent architectures such as BS-Diff [11] and BS-LDM [12] leverage conditional and latent diffusion models for high-resolution bone suppression. Consequently, our contributions establish segmentation as a downstream application on bone-suppressed CXRs, demonstrating its clinical significance.

**Table 1.** Comparison of Bone Suppression Downstream Applications.

| Study | Disease | Dataset | Arch. | Seg. |
| --- | --- | --- | --- | --- |
| Rajaraman et al. [5] | Tuberculosis | Shenzhen, Montgomery TB CXR | VGG-16 | ✗ |
| Rajaraman et al. [6] | COVID-19 | COVID-19 CXR, RSNA | U-Nets and FPNs | ✗ |
| Bae et al. [7] | Lung Nodules | Private | GANs | ✗ |
| Xu et al. [8] | Lung Nodules and Diseases | NODE21, ChestXRy14 | Mask R-CNN | ✗ |
| Han Li et al. [9] | Lung Diseases | Shenzhen, ChestXRy14 | U-Net and CycleGAN | ✗ |
| **This work** | **Pneumothorax** | **SIIM-ACR, PTX-498** | **UNet++, MANet, SegFormer, TransUNet** | **✓** |

## 3. METHODOLOGY

### Datasets

We evaluate the impact of bone suppression on segmentation across two publicly available pixel-level annotated pneumothorax datasets, SIIM-ACR and PTX-498 [13]. SIIMACR, the largest public pneumothorax segmentation dataset consists of 12,047 CXR DICOM files collected from multiple centers under varied acquisition protocols (PA/AP=0.6562; M:F=1.22), resampled to 512 × 512 resolution PTX-498 is a multi-center dataset of 498 pneumothorax-positive CXRs from three hospitals in Shanghai, with pixel-level annotations by two senior radiologists. As shown in Table 2, the SIIM-ACR dataset was upsampled to address class imbalance via random augmentation of positive samples during training. Cross-dataset evaluation enables robust analysis of bone suppression efficacy across varied acquisition protocols and patient distributions.

**Table 2.** Dataset Distribution for Training and Evaluation.

| Phase | Dataset | Positive | Negative | Total |
| --- | --- | --- | --- | --- |
| Original | SIIM-ACR | 2,669 (22.15%) | 9,378 (77.85%) | 12,047 |
| Train | SIIM-ACR | 2,401 (22.15%) | 8,441 (77.85%) | 10,842 |
|  | Upsampled | 8,404 (57.03%) | 6,331 (42.96%) | 14,735 |
| Test | SIIM-ACR | 268 (22.24%) | 937 (77.76%) | 1,205 |
|  | PTX-498 | 498 (100%) | 0 (0%) | 498 |

### Bone Suppression and Pre-processing

Bone suppression was performed using a ResNet-based model [5] trained on 4,500 paired CXRs derived from 247 bone-suppressed JSRT images. The model was optimized using a multiobjective MS-SSIM and MAE loss to minimize structural distortion while preserving soft-tissue cues, achieving low reconstruction error (MS-SSIM = 0.0172, MSE = 0.014). The residual model suppresses bony structures while preserving fine-grained soft-tissue detail, with negligible computational overhead at inference (0.42s/batch). Following suppression, images were standardized and augmented to ensure robustness across varying exposure and acquisition conditions.

### Model Architecture

We benchmark bone suppression across convolutional and transformer-based architectures. UNet++ [14] extends UNet with dense nested skip connections which progressively bridge the semantic gap between encoder and decoder feature maps. While, MANet [1] incorporates Position-wise and Multi-Scale Attention Blocks to capture spatial and channel dependencies across the feature maps. Transformer-based models capture long-range dependencies. SegFormer [15] employs hierarchical multi-scale feature extraction with lightweight MLP decoders, while TransUNet [16] incorporates transformers into the U-Net backbone for combined global and local feature learning.

For feature extraction, InceptionResNetV2 integrates Inception modules with residual connections to capture both fine-grained and high-level features efficiently. R50+ViTB 16 combines ResNet-50’s localized spatial feature extraction with transformer-based global context modeling. Additionally, scSE blocks [17] enhance both spatial and channel-wise feature representations for improved segmentation delineation. As shown in Figure 1, the bone-suppressed input is fed into the encoder backbone for robust feature extraction and then to the decoder blocks of the architecture passing through the segmentation head with a sigmoid activation for mask prediction.

**Fig. 1.**
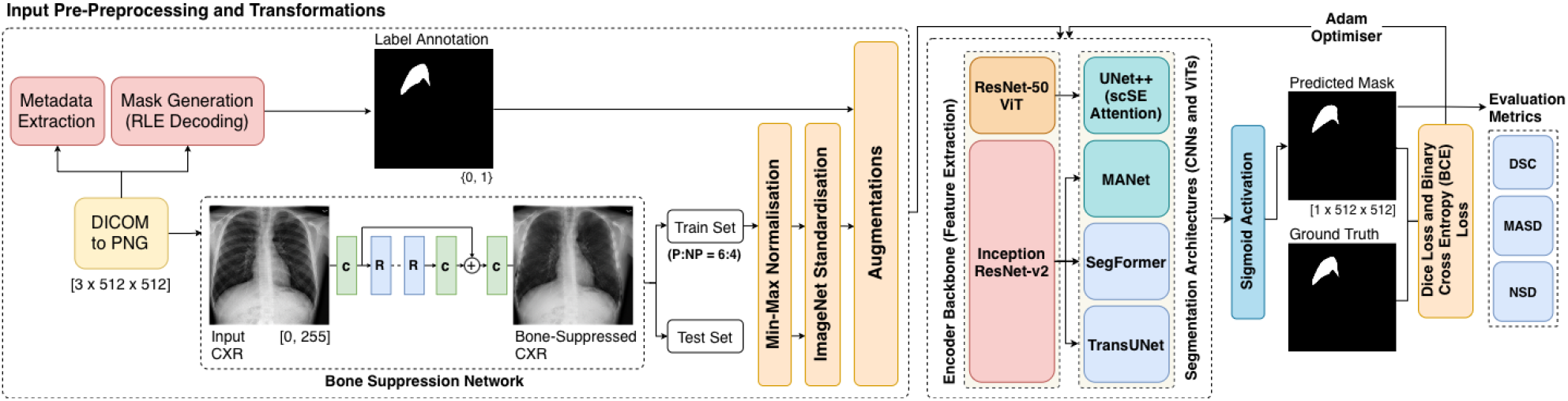
Comprehensive two-stage pipeline for bone suppression and pneumothorax segmentation. Extensive evaluation across two datasets, four segmentation architectures, and three distinct metrics.

### Loss Function

To jointly optimize for global overlap (Dice Loss, *L*_*Dice*_) and pixel-wise accuracy (Binary Cross-Entropy Loss, *L*_*BCE*_), we employed a hybrid loss function defined as: *L*_*Dice*−*BCE*_ = *α*_*D*_ · *L*_*Dice*_ + *α*_*B*_ · *L*_*BCE*_.

### Hyperparameter and Training Strategy

Training was conducted on an NVIDIA RTX A6000 (48GB) for 75-100 epochs with a batch size of 16. Model weights were initialized with ImageNet pre-trained parameters, and Adam optimizer with a learning rate of 0.0001 adjusted via the ReduceLROnPlateau scheduler. A hybrid loss function, combining Dice (*α*_*D*_ = 1) and BCE (*α*_*B*_ = 2) losses, was used to balance global structure preservation and pixel-wise accuracy. Raw logits were passed through a sigmoid activation, with a threshold of 0.5 applied for binary pixel classification

### Evaluation Metrics

Segmentation performance was evaluated using DSC to quantify global overlap, MASD to compute the average distance between predicted and ground truth boundaries, and NSD to measure pixel-level misclassifications near boundaries within a dilated tolerance *τ* [18], thereby ensuring both overall region alignment and precise boundary localization.

## 4. RESULTS

### Bone Suppression Boosts Segmentation

Table 3 shows that training segmentation models on bone-suppressed chest X-rays (BS-CXRs) consistently leads to improved pneumothorax segmentation performance across all architectures. On average, DSC increased by 3.72%, MASD decreased by 10.56%, and NSD improved by 3.89%. The largest individual gains were observed in DSC with 4.89% (MANet), MASD with 17.05% (relative reduction with UNet++), and NSD with 5.88% (MANet), confirming the importance of BS-CXR.

**Table 3.** Performance of Segmentation and Classification Tasks across Architectures on BS-CXR Configurations.

| Architecture | Encoder | BS-CXR | Segmentation |  |  | Classification |  |
| --- | --- | --- | --- | --- | --- | --- | --- |
| | | | DSC (↑) | MASD (↓) | NSD (↑) | F1 (↑) | MCC (↑) |
| UNet++ | Inception-ResNetV2 | – | 0.8064 | 5.4757 | 0.7673 | 0.7695 | 0.7004 |
|  |  | ✓ | **0.8317** | **4.5423** | **0.7903** | **0.7933** | **0.7340** |
| MANet | Inception-ResNetV2 | – | 0.7830 | 5.4116 | 0.7397 | 0.7355 | 0.6569 |
|  |  | ✓ | **0.8213** | **4.8838** | **0.7832** | **0.7788** | **0.7141** |
| SegFormer | Inception-ResNetV2 | – | 0.8088 | 5.3425 | 0.7665 | 0.7577 | 0.6866 |
|  |  | ✓ | **0.8288** | **5.0721** | **0.7882** | **0.8065** | **0.7496** |
| TransUNet | ResNet50-ViT | – | 0.7987 | 5.6073 | 0.7594 | 0.7496 | 0.6744 |
|  |  | ✓ | **0.8342** | **5.0335** | **0.7890** | **0.7973** | **0.7382** |

To quantify the statistical significance of these improvements, we conducted an independent t-test between the results with and without BS-CXRs, using a significance level of *α* = 0.05. The test revealed statistically significant differences in **DSC (p = 0.0018), MASD (p = 0.0025), and NSD (p = 0.0021)**, confirming that the segmentation performance gains on BS-CXRs are architecture agnostic.

Figure 2 provides a visual understanding of the improved localization, boundary delineation, and reduced false positives achieved with bone-suppressed CXR images. These enhancements correspond to: reduced pixel-wise misclassifications at boundaries (NSD and MASD), and better identification of the region of interest (ROI) and localization towards pneumothorax regions (DSC). The GradCAM activation maps illustrate sharper and more focused attention on pathological regions in bone-suppressed CXRs.

**Fig. 2.**
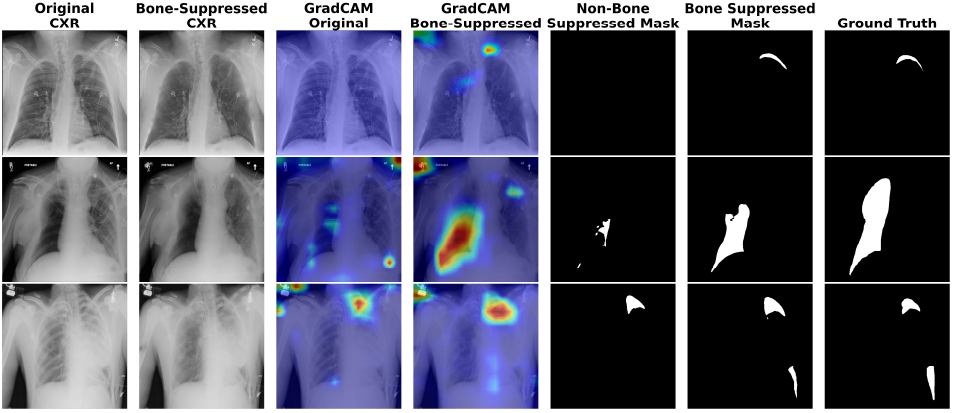
Visual Comparison: BS-CXRs vs. Non-BS-CXRs for MANet (Top), UNet++ (Middle), and SegFormer (Bottom)

### Improvements on Classification

Consistent with the segmentation results, bone suppression also improved pneumothorax classification by enhancing visibility of soft-tissue cues (Table 3). For classification, a case was classified as positive if the model segmented any pneumothorax pixels. Cases with no predicted pixels were predicted negative. F1-scores improved by up to 6.44% (SegFormer). Matthew’s Correlation Coefficient (MCC), a balanced measure of binary classification quality, increased by 9.48% (TransUNet; BS: 0.738, Non-BS: 0.674).

## 5. CONCLUSION

We addressed the challenge of localizing pneumothorax in chest X-rays, where ribs and clavicles obscure collapsed lung regions. We investigated the novel downstream application of bone suppression for segmentation and benchmarked its impact across CNN and ViT-based models on two public datasets, demonstrating statistically-significant architecture agnostic performance gains. Our work advances pleural lung pathology segmentation by integrating bone suppression as a pre-processing step, improving pathological region identification in chest X-rays. Evaluation using boundary-aware metrics quantified improvements in pneumothorax localization accuracy. Future work will explore extending bone suppression to larger-scale datasets and a broader range of applications to improve adaptability and generalization across diverse clinical settings.

## 6. COMPLIANCE WITH ETHICAL STANDARDS

This study utilized publicly available, de-identified datasets (SIIM-ACR and PTX-498) and did not involve any direct patient interaction. Therefore, ethical approval was not required.

## 7. ACKNOWLEDGMENTS

The first three authors (A.S., A.R., and S.S.) are thankful to Harish and Bina Shah School AI & CS and the Office of Research at Plaksha University for providing seed financial support through the Startup Research Grant Ref. No. OOR/PUSRG/2023-24/06 for this research work.

